# Uncovering microbial populations in the lumen of neonatal enteral feeding tubes utilising 16s rRNA sequencing

**DOI:** 10.1101/2020.03.02.972703

**Authors:** Christopher J Winnard, Sue Green, Alison Baylay, Mark J Johnson, Mandy Fader, Charles W Keevil, Sandra Wilks

**Affiliations:** School of Biological Science, University of Southampton, Southampton, UK; School of Health Sciences, University of Southampton, Southampton, UK; Faculty of Health and Social Sciences, Bournemouth University, Bournemouth, UK; Environmental Genomics Facility, University of Southampton, Southampton, UK; Department of Neonatal Medicine, Princess Anne Hospital, University Hospital Southampton NHS Foundation Trust; National Institute for Health Research, Southampton Biomedical Research Centre, University Hospital Southampton NHS Foundation Trust and University of Southampton, Southampton, UK

## Abstract

Gastrointestinal microbiome is increasingly implicated in the morbidity associated with being born preterm. Enteral tubes (ET) are essential for the nutritional care of preterm infants. Limited culture-based studies have suggested they are colonised by high densities of microorganisms. Microbial DNA was extracted from 60 ETs retrieved from infants in a tertiary neonatal unit and analysed by16s rRNA sequencing of the V4 variable region. Relative abundance analysis on dominant microorganisms demonstrated that compared to breast milk, formula significantly increased abundance of *Streptococcus spp* and significantly decreased *Enterococcus spp* and Enterobacteriaceae Vaginal birth was also associated with significantly increased relative abundance of *Streptococcus*. This study more accurately demonstrates the extent of microbial diversity in neonatal ETs, with feeding regime significantly influencing colonisation patterns. Colonisation with unwanted organisms, as a result of specific care regimes, could result in disruption of the fragile infant gut microbiome, with implications for long-term morbidity.

## Background

Despite advances in neonatal medicine that have seen improvements in survival rates of infants born at the extremes of prematurity over the past two decades, there is still a considerable burden of morbidity. In particular, necrotising enterocolitis (NEC), a severe and devastating disease primarily affecting the gastrointestinal (GI) tract of premature infants, inducing inflammation and bacterial invasion of intestinal walls^1^, remains a concern in the preterm population, especially those born with low birth weight. While the precise pathogenesis of NEC is not fully understood, important factors include intestinal ischaemia, gut colonisation by pathologic bacteria, and excess protein substrate in the intestinal lumen^2 3^. Similarly, nosocomial infection remains a significant cause of morbidity and mortality within the extremely preterm population.

Recently there has been increasing interest in the role of host intestinal microbiome in the pathogenesis of NEC and susceptibility to invasive bacterial infection. The hypothesis that the development of a normal commensal flora is fundamental to lifelong health and disease susceptibility is now firmly recognised^4 5^. Perturbations to the maternal, or early infantile, microbiome have been linked to autoimmune and metabolic disorders, such as Type 1 and 2 diabetes^6^. It has also been demonstrated that infants exposed to prolonged courses of antibiotics have increased incidence of NEC, among other severe outcomes^7 8^. This could result from antibiotics hindering or suppressing the development of a ‘normal’ microbial flora, instead allowing for selection of more resistant pathogenic bacteria, increasing risk of morbidity and mortality. In addition, routine use of probiotics in an attempt to modify bacterial colonisation in preterm infants has been shown to significantly reduce the risk NEC and improve feed tolerance^9^.

Preterm infants are exposed to a variety of factors that result in them having abnormal intestinal flora compared to infants born at term; they are more likely to be born by caesarean section, are often treated with board spectrum antibiotics and spend prolonged periods in a hospital environment where they are exposed to a variety of nosocomial organisms. In addition, they often have indwelling medical devices such as central venous catheters and enteral feeding tubes, which further influence their exposure to microorganisms. Compared to term-born infants, preterm infants have abnormal faecal colonisation, with a paucity of normal enteric bacterial species, and delayed bacterial colonization^10 11^. Feeding with maternal breast milk (MBM) is the most effective preventative mechanism against development of NEC^12 13^, and one reason for this may be that it promotes the growth of non-pathogenic bacterial species while reducing the pathogen load^14^. MBM has also been shown to block bacterial lectins^15^, significantly reduce the risk of infection^16^, improve feed tolerance and have longer term benefits on growth and neurodevelopment^17^. Many of these effects may also be mediated through alterations in intestinal bacterial flora. Whilst MBM is beneficial, it is not always available and so alternative feeds must be utilised; with many studies demonstrating that these alternative feeds can impact microbiome development^18 19^. Feed type therefore clearly has implications for microbiome development and outcomes in preterm infants.

Early research into the understanding of the microbiome relied mainly on culture-based analysis of microbes; however, more recently developed techniques are revealing that culture-based analysis alone has led to a severe underrepresentation^20^. A growing wealth of literature suggests that, for many bodily and environmental sites, the greater proportion of microbes present may in fact be unculturable by traditional recovery techniques. A study using molecular based methods to gain an understanding into the human intestinal microbial flora demonstrated that 80% of the microbes uncovered were unculturable, with 60% being novel at the time^21^. 16s rRNA sequencing is rapidly providing more accurate representations of microorganisms within a given environment. Such methods are also able to identify species that may not be recoverable by culture-based analysis, such as viable but non-culturable (VBNC) species.

Previous studies have analysed alpha (diversity of microbial communities within one site) and beta (dissimilarities in microbial diversity between two sites) diversity of faecal samples to assess impact of common infant characteristics and care regimes (feed types, antibiotic prescription etc). Given the links between feed type, bacterial colonisation and outcomes in preterm infants, we sought to accurately portray the microbial communities found within neonatal feeding tubes, utilising modern molecular analysis techniques, as well as gain insight into how care practices in early life influence their dynamics. This novel approach could potentially lead to more personalised medicine, providing vital information to clinicians and enabling tailoring of care practices to differently vulnerable infant populations.

## Methods

### Ethics

Ethical approval was obtained (NHS Research and Ethics Committee: East of Scotland Research Ethics service, Reference 17/ES/0142). As per the ethical approval, informed consent for tube collection was not required as samples analysed were those designated for disposal, no human genetic material was collected and no patient identifiable information was collected, stored or analysed.

### Setting and population

All tubes were collected from a tertiary Neonatal Intensive Care unit offering surgical and other speciality services. All nasogastric or orogastric feeding tubes removed from infants within the unit as part of their normal care regime between April and June 2018 were included unless they were damaged upon removal/collection or were removed within an emergency situation where there was inadequate time for appropriate sample collection. Clinical data collected included date of tube removal, tube characteristics (including length, diameter and material type), duration of placement, reason for removal, type of tube (nasogastric or orogastric), gender of infant, gestational age (GA) of infant at birth, GA at tube removal, mode of delivery, feed type, nutritional supplement use, antibiotic exposure and mode of ventilation. Feed type was categorised into MBM, donor breast milk (DBM), infant formula (IF) and nil by mouth (NBM). GA at birth and at time of tube removal was categorised according to World Health Organisation definitions as extremely preterm (< 28 weeks), very preterm (28 to 32 weeks), moderately preterm (32 to 37 weeks) and term (> 37 weeks).

### Sample processing and DNA extraction

After removal from infants, feeding tubes were placed directly into sterile bags, sealed and stored at 4°C within the unit’s designated research refrigerator preceding collection. Samples were collected and transported to a containment level 2 category laboratory. After sterilisation of the external surfaces of the tubes, sterile Dulbecco’s Phosphate-buffered saline (DPBS) was flushed through the tubes and collected. Tubes were cut into approx. 1 cm segments. Segments were vortexed within tube flush to dislodge all microbes. Cell suspension was removed and centrifuged to pellet cells. Genomic DNA was isolated from entire pellet with the QIAmp Ultraclean production (UCP) pathogenic mini kit (Qiagen, Germany) following manufacturers guidelines with the following modification for optimal DNA extraction: QIAmp pathogen lysis tubes (L) for mechanical lysis in a TissueLyser LT for 15 mins at 50hz, as outlined in Pre-treatment of Microbial DNA protocol, before DNA extraction following manufacturers Sample Prep (Spin protocol) guidelines. Total DNA eluted into 30µl buffer AVW (RNase-free water with 0.04% NaN3 (sodium azide)) and yield measured by NanoDrop™ spectrophotometer before being stored at −80°C. Once 60 samples had been reached, they were transferred to an external company (Environmental Genomics Facility, University of Southampton, UK) for 16s rRNA sequencing.

### Analysis of amplicon sequencing data

The V4 region of the 16S rRNA gene was amplified from all 60 samples using fusion primers 515F:*TCGTCGGCAGCGTCAGATGTGTATAAGAGACAG*GTGYCAGCMGCCGCGGTAA and 806R:

*GTCTCGTGGGCTCGGAGATGTGTATAAGAGACAG*GGACTACNVGGGTWTCTAAT, which consist of the V4 region primers^22^ ligated to Illumina Nextera adaptor consensus sequences (indicated in italics). PCR reactions were carried out in 25 µl volumes, consisting of 12.5 µl NEBnext Q5 HiFi Hotstart mastermix, 12.5 ng genomic DNA, and 1 µl forward and reverse primers (10 µM), and amplified using the following conditions: 95°C for 3 min, followed by 25 cycles of 95°C for 30 sec, 55°C for 30 sec and 72°C for 30 sec, with a final extension of 7 min at 72°C. Amplicons were cleaned using 0.8 x volume AMPure XP beads (Beckman Coulter Ltd, UK) and dual indexed using a Nextera XT v2 Index Kit (Illumina, United States) using a further 8 PCR cycles. The resulting amplicon libraries were pooled and sequenced on an Illumina MiSeq instrument, using a MiSeq v3 Reagent Kit (Illumina) and 2 × 300 bp paired end sequencing. Sample demultiplexing was carried out on-instrument by the MiSeq control software.

Qualitative Insights into Microbial Ecology 2 (QIIME2 version 2018.8) was used for analysis of 16s rRNA gene amplicon sequences. Demultiplexed reads were trimmed using Cutadapt version 1.17 ^23^ to remove residual adapters and primers, and to remove low-quality 3’ bases (quality threshold 20). Reads less than 250 bp following trimming were discarded. Denoising was carried out using the DADA2^24^ plugin within QIIME2. Taxonomy was assigned to the resulting amplicon sequence variants (ASVs) using the naïve-Bayes machine learning classifier method implemented in QIIME2’s q2-feature-classifier plugin^25^ (taxonomic assignment against Greengenes 13_8 99% classifier). Keemei plugin was utilized for metadata file validation^26^.

Diversity analysis included both alpha and beta diversity. Alpha diversity was assessed using both species richness (Faith’s Phylogenetic Diversity (PD)) and community evenness (Pileu’s evenness), combined with Kruskal-Wallis total and pairwise analysis. Beta diversity (extent of change in community composition) was assessed using non-parametric multivariate PERMANOVA pairwise analysis. QIIME biom convert function provided summary information of relative abundance of taxonomic groups for each taxonomic level. Analysis of relative abundance of main bacterial genus: *Enterobacteriaceae, Staphylococcus, Streptococcus, Enterococcus* and *Neisseria*, grouped relative to patient information was conducted. Data was tested for normality using Shapiro-Wilk tests. For non-normally distrusted data, Kruskal-Wallis H tests were conducted for each. For groups with significant differences Dunn’s post hoc pairwise multiple comparisons were conducted. Significance was set at p < 0.05.

## Results

### Infant characteristics

Infants were included in the study with a mean (standard deviation, SD) GA at birth of 29.86 (5.14) weeks. 60 tubes were collected from 30 infants throughout the study with a mean (SD) of 2 (1.2) tubes per infant and range 1-6 tubes. For summary statistics refer to Table 1. Predominant characteristics of the samples were: male, born extremely premature although within the moderately premature category at tube removal, not receiving antibiotics, ventilated, receiving vitamin or iron supplements but receiving MBM, with a 6 french gauge nasogastric tube placed for 7 days and routinely removed.) For further patient information refer to supplementary materials Figure S1.

**Table 1.** Summary patient information for all 60 samples included within study.

| Variable | n (n=60) | % |
| --- | --- | --- |
| Sex |  |  |
| F | 25 | 42% |
| M | 35 | 58% |
| GA at Birth* |  |  |
| Extremely Preterm | 24 | 40% |
| Very Preterm | 21 | 35% |
| Moderately Preterm | 6 | 10% |
| Term | 9 | 15% |
| GA at Removal |  |  |
| Extremely Preterm | 3 | 5% |
| Very Preterm | 13 | 22% |
| Moderately Preterm | 29 | 48% |
| Term | 15 | 25% |
| Route of Birth |  |  |
| Vaginal | 36 | 60% |
| Ceaserian Section | 24 | 40% |
| Antibiotic Prescription |  |  |
| Y | 18 | 30% |
| N | 42 | 70% |
| Vitamin Supplement |  |  |
| Y | 24 | 40% |
| N | 36 | 60% |
| Iron Supplement |  |  |
| Y | 12 | 20% |
| N | 48 | 80% |
| Overall Ventilation |  |  |
| Y | 18 | 30% |
| N | 42 | 70% |
| Ventilation Type |  |  |
| Invasive | 9 | 15% |
| Non-Invasive | 9 | 15% |
| None | 42 | 70% |

\*GA classed according to WHO: <28 weeks = Extremely preterm. 28-32 weeks = Very Preterm. 32-37 week = Moderate to Late preterm. >37 weeks = Term.
\*\*Invasive types: SIMV (4), PC-SIMV (5), PC-AC (15), PC-AC + VG (3), PC-AC + Nitric (1), Optiflow (1). Non-Invasive: CPAP (14), Bubble-CPAP (4), Nippy-CPAP (1), High-Flow (4 and Vapotherm (6)). None: All infants not receiving ventilation, Low-flow O2 (3) and Nasopharyngeal Airway (1).
Abbreviations: GA : Gestational Age. SIMV : Synchronized Intermittent-Mandatory
Ventilation. PC : Pressure Controlled. AC : Assist Controlled. CPAP : Continuous Positive Airway Pressure. VG : Volume Guaranteed.

### Common bacterial phyla

The most dominant phylum throughout the entire sample population was *Firmicutes*, followed closely by *Proteobacteria. Actinobacteria* also displayed prominently in many samples. Nearly all the *Firmicutes* present were *Bacilli* - more specifically of the order *Lactobacillales* and *Bacillales* with minor populations of *Gemellales* and *Clostridiales* present. The *Proteobacteria* populations were comprised of predominantly *Gammaproteobacteria* followed by *Betaproteobacteria* and within these *Enterobacteriales* and *Neisseriales* respectively.

### Alpha diversity analysis

The Alpha-group-significance command was used to analyse microbial compositions in relation to sample metadata. Faith’s PD analysis of community richness was measured for all categorical patient variables, demonstrating that categorical variables associated with overall significant differences in community richness were: GA category at birth (p = 0.007), mode of delivery (p = 0.021), where infants born via caesarean section displayed a lower community richness that those born vaginally, and feed type (p = 0.049).

Pairwise analysis (Kruskal-Wallis) of diversity comparing GA categories revealed significant differences between extremely preterm infants compared to very preterm infants (p = 0.002), and term infants compared to very preterm (p = 0.009). Extremely preterm infants had the highest microbial richness (Faith’s PD value), whilst very preterm infants had the lowest. For main feed type, although there was overall group significance, pairwise analysis did not detect significant differences between subgroups. Infants fed DBM had the highest average community richness, while being fed IF resulted in lowest species richness.

Pileu’s evenness (measure of community evenness) was then calculated. Variables associated with significant differences in evenness were GA category at removal (p < 0.001), mode of delivery (p = 0.023), antibiotic exposure (p = 0.033) and vitamin usage (p = 0.005). Similar to community richness, those born vaginally also displayed higher community evenness than those born via caesarean section. Those receiving antibiotics had decreased evenness compared to those not receiving treatment, whereas those who were receiving vitamin supplements had increased community evenness.

### Beta Diversity

PERMANOVA analysis was utilised to test the distances between samples from within a subgroup in relation to clinical data, in order to ascertain if samples from within categorical subgroups were more similar to one another or to the other subgroups. Clinical variables found to have significant differences between their subgroups were; main feed type (p = 0.002), GA at birth (p = 0.005), GA at tube removal (p = 0.007), antibiotic exposure (p = 0.013) and gender (p = 0.038), though mode of delivery was not significant (p = 0.06). Categorical variables with more than two subgroups were analysed against one another using pairwise PERMANOVA permutation tests. For main feed type all feed subgroups were found to be significantly different from one another: DBM versus IF, MBM and NBM (p = 0.003, 0.002 and 0.042 respectively), IF versus MBM and NBM (p = 0.050 and 0.029 respectively) and MBM versus NBM (p = 0.015). For GA at birth, there was a significant difference seen in extremely preterm infants compared to very preterm (p = 0.007). For GA at tube removal, there were significant differences between extremely preterm compared to both moderately preterm and term infants (p = 0.013 and 0.039 respectively), and for moderately preterm versus term (p = 0.013).

Multiple multivariate response linear regressions were run utilising an increasing amount of patient variables. Summary r^2^ values, indicating the percentage of community variation the regression model can explain, increased with increasing numbers of patient variables included, however this also decreased the r^2^ values of each individual covariate. Some over fitting occurred in the expanded model as the fold 0-cross validation prediction accuracy was higher than within model error, however, all other cross folds, and all within the reduced model, were lower.

Summary r^2^ values for expanded and reduced models were 0.4792 and 0.3249 respectively, suggesting that even the expanded model only accounted for around 48% of community variation. For the expanded model the most influential variable was a GA category at birth of term, followed closely by a GA category at tube removal of term. Whereas for the reduced model, it was a GA category at birth of moderately preterm, followed closely by term. In the expanded model all variable categories appear to be responsible for variation of around 1-2%, and this is similar in the reduced model at around 1-3%.

### Relative abundance of dominant phyla

Visual inspection of Figure 1, displaying relative frequency of species at a higher taxonomic (Class) level split based on main feed type, provides some insight into the significant differences observed. Analysis of relative abundance of main bacterial family or genus: *Enterobacteriaceae, Staphylococcus, Streptococcus, Enterococcus* and *Neisseria*, grouped relative to feed type was conducted. Shapiro-Wilk normality tests suggested data was not normally distributed and so Kruskal-Wallis H tests were conducted for each. For groups with significant differences Dunn’s post hoc pairwise multiple comparisons were conducted with confidence levels set at p < 0.05. Refer to Table 2, for relative abundance data summary.

**Table 2.** Mean, standard deviation (SD) and range, for relative abundance of main microorganism; *Staphylococcus, Enterococcus, Streptococcus, Neisseria* and *Enterobacteriaceae*. Relative abundance of total microorganisms is capped at 1, where 1 is complete dominance of the entire sample. Samples with relative abundance 0 (absent from sample) of a given microorganism were removed from results for this table.

| Microorganism | Relative Abundance |  |
| --- | --- | --- |
|  | Mean (SD) | Range |
| *Staphylococcus* | 0.272 (0.272) | 0.00044 to 0.991 |
| *Enterococcus* | 0.136 (0.203) | 0.00005 to 0.771 |
| *Streptococcus* | 0.148 (0.211) | 0.0002 to 0.715 |
| *Neisseria* | 0.085 (0.145) | 0.00004 to 0.668 |
| *Enterobacteriaceae* | 0.221 (0.284) | 0.00003 to 0.931 |

**Figure 1.**
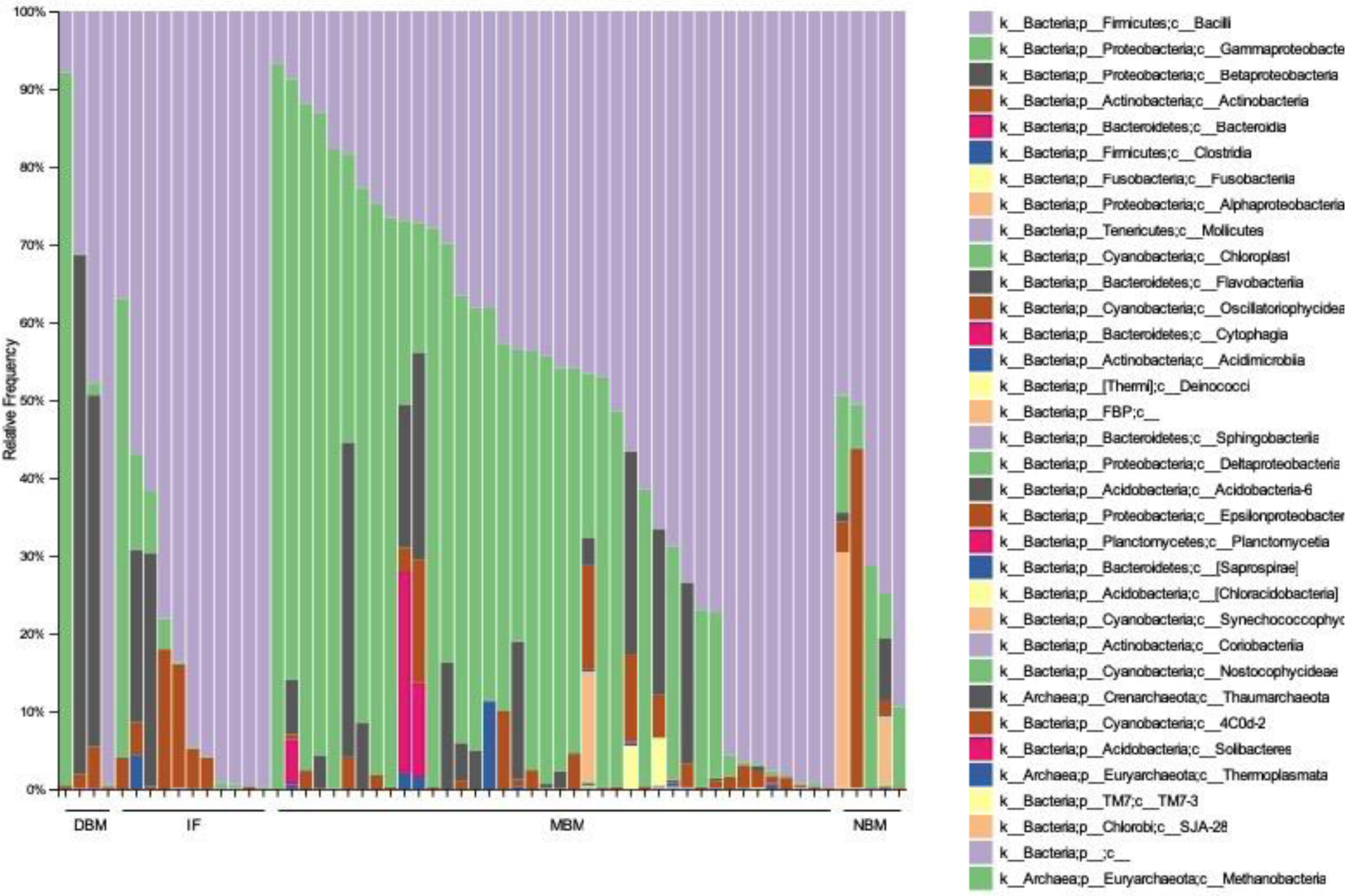
Taxonomic plot displaying relative frequency for all 60 samples split based on main feed category, where; DBM= donor breast milk, IF= infant formula, MBM= maternal breast milk and NBM= nil by mouth.

Overall significant differences between feeding groups was found for relative abundance of *Enterococcus* (p = 0.002), *Streptococcus* (p < 0.001) and *Enterobacteriaceae* (p = 0.004). Dunn’s multiple comparison test was used to distinguish statistically significant differences between feed subgroups. For *Streptococcus*, the IF subgroup was found to have significantly higher relative abundance than the DBM (p < 0.001) and MBM (p < 0.001). Whereas for *Enterococcus*, the IF subgroup was found to have significantly lower relative abundance from all other feed subgroups (p < 0.04 for all), as can been seen in Figure 2. For *Enterobacteriaceae* only the IF and MBM subgroups were found to be statistically significantly different (p = 0.003), with MBM being significantly higher. There were no significance differences in relative abundance between feeding groups for *Staphylococcus* or *Neisseria*.

**Figure 2.**
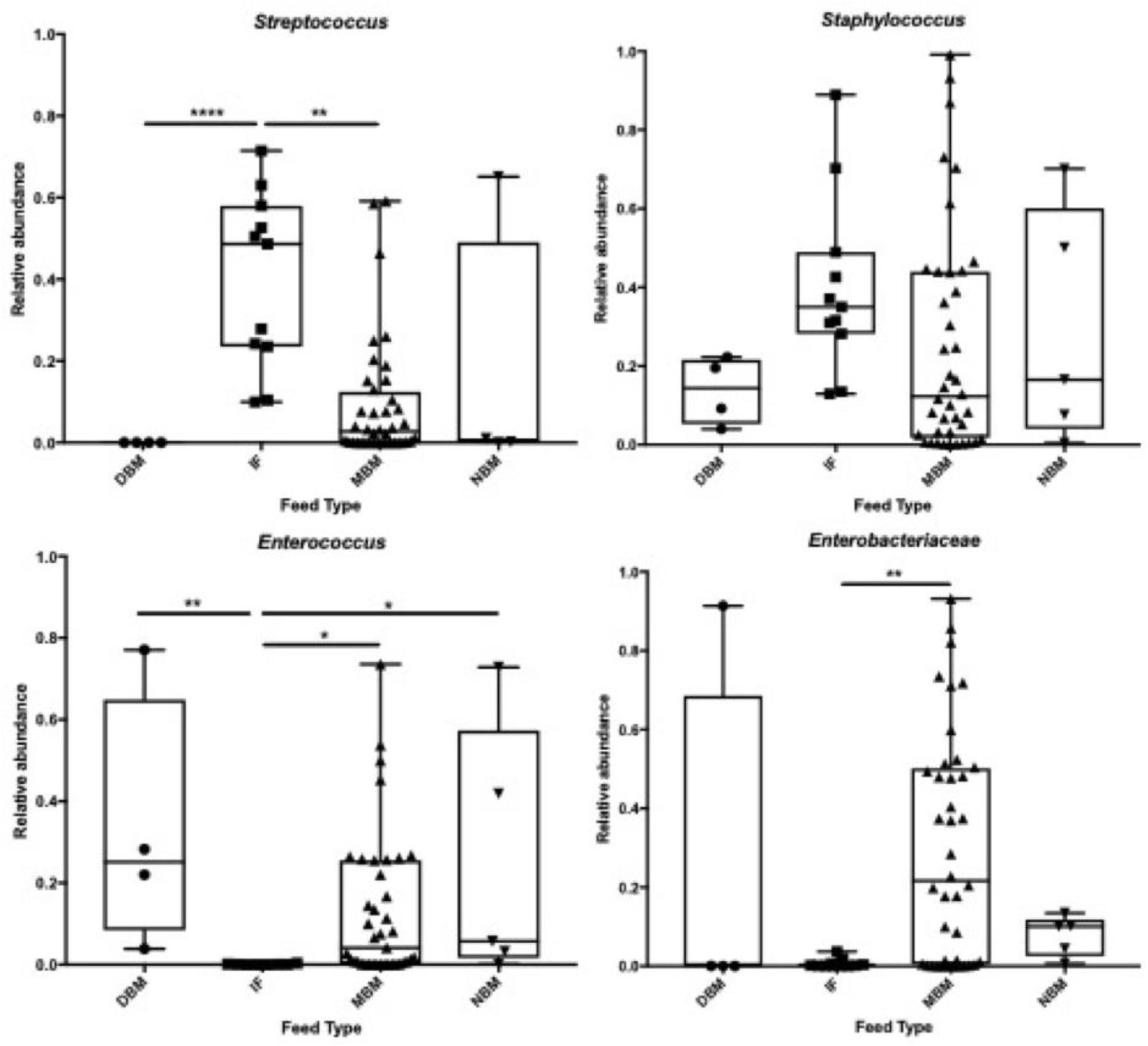
Box plots for relative abundance classified to family level for 4 most abundant profiles; *Streptococcus, Staphylococcus, Enterococcus* and *Enterobacteriaceae*, against main feed type. Relative abundance of total microorganisms is capped at 1, where 1 is complete dominance of the entire sample. Samples with relative abundance 0 (absent from sample) of a given microorganism were not removed from results. p-values as indicated by ns = p > 0.05, * = p ≤ 0.05, ** = p ≤ 0.01, *** = p ≤ 0.001 and **** = p ≤ 0.0001.

Analysis of main family of phylum was conducted for other patient characteristics. *Streptococcus* relative abundance plotted against mode of delivery (Figure 3) was found to be statistically significant (p = 0.037).

**Figure 3.**
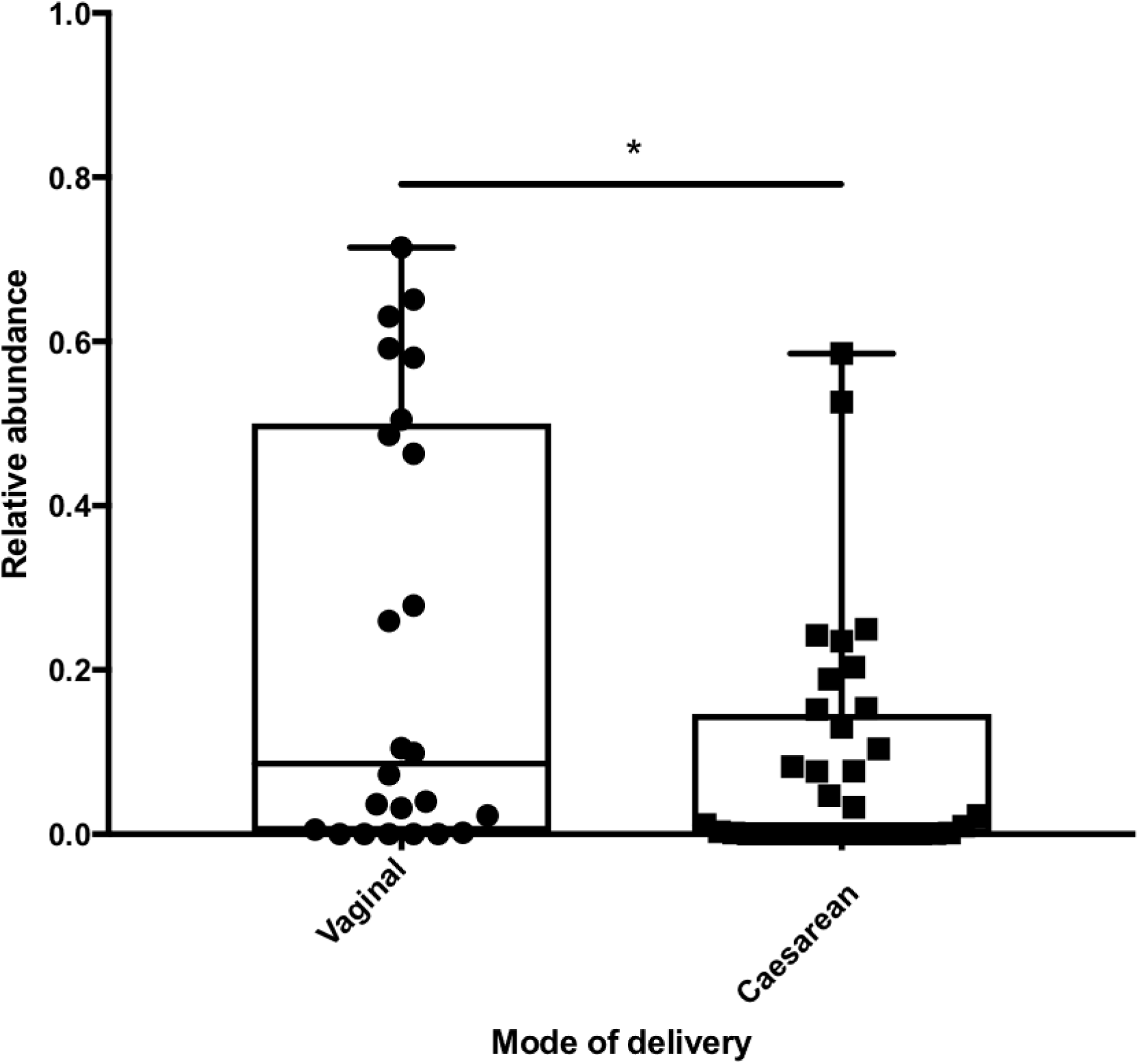
Displays relative abundance of *Streptococcus* split by mode of delivery. p-values as indicated by ns = p > 0.05, * = p ≤ 0.05, ** = p ≤ 0.01, *** = p ≤ 0.001 and **** = p ≤ 0.0001.

The analysis of both GA categories at birth and tube removal revealed only relative abundance of *Staphylococcus* against GA at birth (p = 0.020) was statistically significant. Dunn’s multiple comparison analysis demonstrated that it was the extremely preterm and term categories that differed significantly (p = 0.012). From Figure 4 it can be seen that relative abundance of *Staphylococcus* increased with increasing GA at birth. Although not significant (p = 0.914), GA at tube removal also appeared to follow the same trend. Other patient characteristics (e.g. antibiotic exposure or vitamin usage) were not found to significantly impact relative abundance when plotted individually.

**Figure 4.**
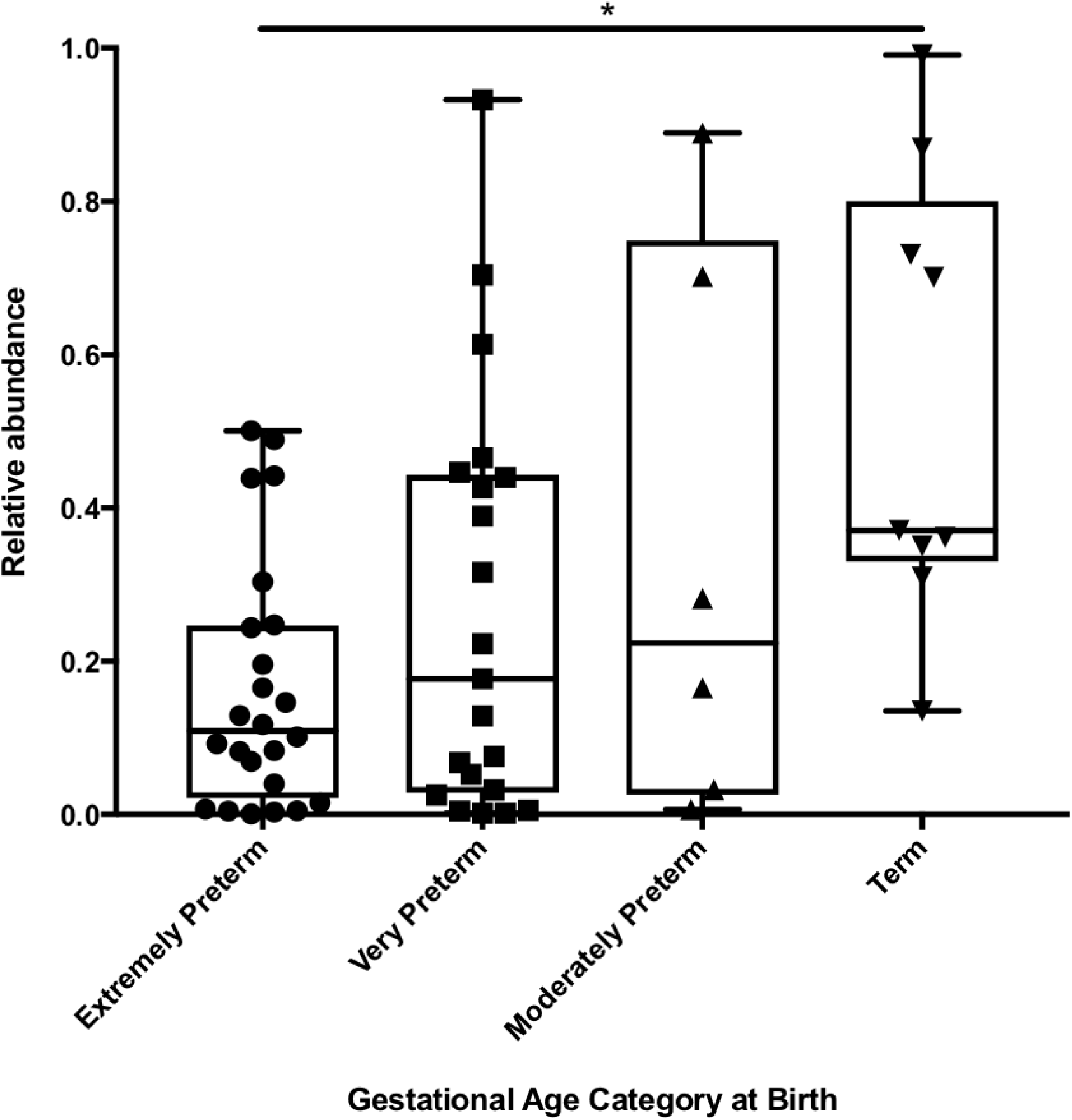
Graph displaying relative abundance of *Staphylococcus* against gestational age category at birth. Relative abundance capped a 1, where 1 is complete sample dominance and 0 is absence of sequences from sample. p-values as indicated by ns = p > 0.05, * = p ≤ 0.05, ** = p ≤ 0.01, *** = p ≤ 0.001 and **** = p ≤ 0.0001.

## Discussion

### Diversity analysis

Previous research has suggested links between levels of microbial diversity and human health and disease^27^. Alpha diversity analysis results from the study reported here demonstrates the significant influence of mode of delivery on microbial communities found within the feeding tubes. This is in line with previous work, utilising infant faeces, demonstrating that vaginally born infants have a greater abundance of species, where their microbiota is seeded by the vaginal and faecal flora of the mother, whereas caesarean born infants microbiota is more influenced by the local environment and the mother skin microbiota^28^. Another dominant influencer on alpha diversity was GA category at birth, and at tube removal. This is in keeping with previous observations that the microbiome of many bodily sites matures over the first few months of life, with preterm infants displaying altered development and communities^29 30^.

Beta diversity analysis demonstrated the pivotal impact of feeding regime upon the maturation of specific microbial communities, a known influence on the neonatal gut microbiome, as analysed through faecal samples^31^. However, it has been suggested that faecal samples underrepresent the biodiversity of the gastrointestinal system^32^. Others have analysed stomach contents, aspirated through feeding tubes^33^, but without knowing the exact contents of the feeding device; this also introduced bias. The placement of enteral feeding tubes directly into the stomach, provides researchers analysing them a unique insight into the dynamics of the gut microbiome, without the influence of other bodily sites. The contents of neonatal feeding tubes provide a truly representative sample of the neonatal gut microbiome, as a result of aspiration checks, prior to each feeding, collecting microorganisms from within this niche and depositing them upon an abiotic surface, away from the influence of the host immune system.

### Abundance analysis

Abundance analysis of the main microorganisms’ present demonstrated how the tubes that were exposed to infant formula had altered populations, with increased relative abundance of *Streptococcus*, but decreased abundance of *Enterococcus* and *Enterobacteriaceae*. Those born vaginally also displayed significantly increased relative abundance of *Streptococcus* within the tubes compared to those delivered by caesarean section. When combined these suggest that an initial increased risk of seeding by *Streptococcus* during vaginal birth could be exacerbated by *Streptococcus* overgrowth upon infant formula feeding, as well as suppression of potentially beneficial populations. This is especially concerning given the association of *Streptococcus* with neonatal early-onset sepsis and meningitis^34 35^. Conversely, infants displaying early colonisation with *Enterococcus* are less likely to develop allergy later in life^36^, perhaps through suppression of inflammatory responses^37^. Larger cohort-controlled studies should be conducted to assess the specific impact of vaginal birth and subsequent formula feeding as a risk factor for *Streptococcus* related morbidity and mortality within the neonatal population. This could lead to the creation of specific prophylactic care regimes or vigilance schemes, aimed at reducing risk within a subpopulation.

### Implications

Despite the relative lack of knowledge on the pathogenesis of NEC, it is clear that bacteria play, at least, a contributory role^38 39^. Bacterial lipopolysaccharide is able to induce activation of TLR-4 and therefore stimulate the inflammatory environment associated with NEC^40^. Specific populations of bacteria may also dominate the gastrointestinal environment and secrete toxic metabolites and endotoxins, worsening inflammation and increasing the risk of potential intestinal damage. Research into specific populations of bacteria associated with NEC have yielded contrasting results, with many identifying bacteria often found in healthy controls^41-43^. These studies have led some to conclude that it may not be one specific pathogenic species contributing, but instead the culmination of many species distorting the normal microbial environment^44^. Colonisation of enteral feeding tubes, as a result of early infant seeding with potentially pathogenic microorganisms, translocation up the tubes as a result of aspiration, or introduction via contaminated external sources, could result in unwanted pathogenic overgrowth within this important niche. Due to the direct placement of these tubes into the infant stomach, any pathogens would be able to not only continually seed their fragile microbiome, leading to dysbiosis, but could potentially invade the host tissue following any device related damage to the gut mucosa.

Given the variability in causative organisms for neonatal infection^45^, investigations into potential sources of infection should utilise techniques that can accurately portray all microorganisms. Previous studies investigating contamination have relied on limited culture-based analysis^46^. As previously mentioned, culture-based analysis alone can lead to severe underrepresentation of microbial diversity, which in a clinical scenario could lead to inaccurate targeted therapies. Despite their limitations, culture analysis of contaminated enteral feeding tubes still demonstrate that high levels of pathogenic colonisation frequently occur^47^. The use of modern molecular techniques to fully describe the microbial populations in neonatal enteral feeding tubes is a particular strength of the present study. Increasingly, recent molecular based analysis studies are showing the importance of minor microbial populations on microbiota formation^48^, and future work should focus on utilising the most accurate methods.

### Limitations

Although a substantial study, sample numbers meant we were not able to adequately control for all cofounders. This was highlighted by the multiple multivariate response linear regression analysis in which all infant characteristics were shown to contribute to community variation to a similar degree. This was most likely a result of the sample size splitting certain populations into very minor subgroups.

## Conclusions

The results demonstrate that early nutrition significantly influences colonisation patterns, and that a combination of vaginal birth and subsequent infant formula feeding may be a specific risk factor for *streptococcal* contamination of enteral feeding tubes. To further elucidate the impacts of early life feeding and care regimes impact microbiome development, a large-scale observational study should be conducted to analyse the impact of infant formulae feeding on vaginally born infants’ infection and morbidity, specifically those attributed to *Streptococcus*. Future studies should also be conducted utilising larger cohort sizes to more accurately account to cofounding patient variables. The therapeutic utilisation of probiotic microorganisms to modify early stage colonisation, as well as restrict potentially pathogenic microorganism overgrowth, of formula fed infants, to more closely resemble those fed breast milk, also provides an exciting avenue for future research on tube colonisation. Overall this study demonstrates the feasibility of 16s rRNA sequencing as a tool for uncovering previously undiscovered microbial populations within the neonatal population.

## Acknowledgements

Funding for the study was provided by University of Southampton, Biological Sciences, Institute for Life Sciences and Health Sciences departments as part of a studentship awarded to CJW. We would like to acknowledge the help of Dr Edward Andrews in clinical data collection. In addition, we would like to thank Jenny Pond, Christie Mellish and Philippa Crowley, neonatal research nurses, as well as all the clinical nursing staff on Southampton Neonatal unit for their help with sample collection

This study was supported by the National Institute for Health Research through the NIHR Southampton Biomedical Research Centre and Southampton Welcome Trust Clinical Research Facility specialist neonatal research nurses for sample collection and site management. MJJ is supported by the National Institute for Health Research through the NIHR Southampton Biomedical Research Centre.

